# A sparse EEG-informed fMRI model for hybrid EEG-fMRI neurofeedback prediction

**DOI:** 10.1101/599589

**Authors:** Claire Cury, Pierre Maurel, Rémi Gribonval, Christian Barillot

## Abstract

Measures of brain activity through functional magnetic resonance imaging (fMRI) or Electroencephalography (EEG), two complementary modalities, are ground solutions in the context of neuro-feedback (NF) mechanisms for brain rehabilitation protocols. While NF-EEG (real-time neurofeedback scores computed from EEG signals) have been explored for a very long time, NF-fMRI (real-time neurofeedback scores computed from fMRI signals) appeared more recently and provides more robust results and more specific brain training. Using simultaneously fMRI and EEG for bi-modal neurofeedback sessions (NF-EEG-fMRI, real-time neurofeedback scores computed from fMRI and EEG) is very promising to devise brain rehabilitation protocols. However, fMRI is cumbersome and more exhausting for patients. The original contribution of this paper concerns the prediction of bi-modal NF scores from EEG recordings only, using a training phase where EEG signals as well as the NF-EEG and NF-fMRI scores are available. We propose a sparse regression model able to exploit EEG only to predict NF-fMRI or NF-EEG-fMRI in motor imagery tasks. We compared different NF-predictors steaming from the proposed model. We showed that predicting NF-fMRI scores from EEG signals adds information to NF-EEG scores and significantly improve the correlation with bi-modal NF sessions, compared to classical NF-EEG scores.

## 1 INTRODUCTION

Neurofeedback approaches (NF) provide real-time feedback to a subject about its brain activity and help him or her perform a given task (Hammond, 2011; Sulzer et al., 2013). The estimation of neurofeedback information, is done through online brain functional feature extraction, to provide this real-time feedback to the subject. NF appears to be an interesting approach for clinical purposes, for example in the context of rehabilitation and psychiatric disorders (Sulzer et al., 2013; Birbaumer et al., 2009; Wang et al., 2017). Functional magnetic resonance imaging (fMRI) and electro-encephalography (EEG) are the most used noninvasive functional brain imaging modalities in neurofeedback. EEG measures the electrical activity of the brain through electrodes located on the scalp. EEG has an excellent temporal resolution (milliseconds), but a limited spatial resolution (centimeters) implying a lack of specificity. Furthermore, source localisation in EEG is a well-known ill-posed inverse problem (Grech et al., 2008).

On the other hand, blood oxygenation level dependent (BOLD) fMRI, measures a delayed hemodynamic response to neural activity with a good spatial resolution, and a temporal resolution of 1 or 2 seconds depending on the sequence used. Therefore fMRI is more specific than EEG, making the fMRI an adequate modality for neurofeedback (NF-fMRI) (Thibault et al., 2018). However the use of the MRI scanner is costly, exhausting for patients since staying perfectly still when suffering is challenging and time consuming, hence NF-fMRI sessions cannot be repeated too many times for the same subject or patient.

During the past few years, the use of simultaneous EEG-fMRI recording has been used to understand the links between EEG and fMRI in different states of the brain activity and received recognition as a promising multi-modal measurement of the brain activity (Perronnet et al., 2018; Abreu et al., 2018). However this bi-modal acquisition is cumbersome for subjects or patients, due to the use of the fMRI scanner. The methodology to extract information from fMRI with EEG have been also intensively investigated (some methods involved in the process are reviewed here (Abreu et al., 2018)). Indeed, both modalities are sensitive to different aspect of brain activity, with different speeds. EEG provides in real time a direct measure of the changes in electrical potential occurring in the brain, while fMRI indirectly estimates brain activity by measuring changes in BOLD signal reflecting neuro-vascular activity, which occurs in general few seconds after a neural event (Friston et al., 1994; Logothetis et al., 2001). Several studies have investigated correlations between EEG signal and BOLD activity, in specific and simple tasks (de Munck et al., 2007; Goncalves et al., 2008; Engell et al., 2012; Magri et al., 2012; Scheeringa et al., 2011), and found different relationships between certain frequency bands on the EEG signal with the BOLD signal. All those studies reveal the existence of a link between EEG and fMRI, but this relationship highly varies with the task, the location in the brain and the considered frequency bands.

In the literature, the term EEG-informed fMRI refers to methods extracting features from EEG signals in order to derive a predictor of the associated BOLD signal in the region of interest under study. A recent review (Abreu et al., 2018) gives a good overview of the principal EEG-informed fMRI methods and their limitations. Different strategies have been investigated, depending on the type of activity under study (epilepsy, resting state, open/closed eyes, relaxation): either by selecting one channel on interest, either by using multiple channels, before extracting features of interest. For example in (Leite et al., 2013; Formaggio et al., 2011), authors used a temporal independent component analysis to select the best channel reflecting the epileptic seizures. In (Schwab et al., 2015), authors used a spatial, spectral and temporal decomposition of the EEG signals to map EEG on BOLD signal changes in the thalamus. From a more symmetrical way, in (Noorzadeh et al., 2017) it has been proposed a method for the estimation of brain source activation, improving its spatio-temporal resolution, compared to EEG or BOLD fMRI only. However, in the context of neurofeedback, using simultaneous recording of EEG-fMRI to estimate neurofeedback scores computed in real time from features of both modalities (NF-EEG-fMRI) is a recent application that have been first introduced, and its feasibility demonstrated by (Zotev et al., 2014; Perronnet et al., 2017; Mano et al., 2017). The recent methodology synchronising both signals for real time neurofeedback (Mano et al., 2017), allows the setup of a new kind of data named NF-EEG-fMRI data, such as the dataset presented by (Perronnet et al., 2017), which we used for the present study. Furthermore, it has been shown in (Perronnet et al., 2017), that the quality of neurofeedback session is improved when using simultaneously both modalities, in NF-EEG-fMRI sessions. Thus, being able to reproduce in real time a NF-EEG-fMRI session when using EEG only, would reduce the use of fMRI in neurofeedback, while increasing the quality of NF-EEG sessions. To export fMRI information outside the scanner, most of the methods intend to predict the fMRI BOLD signal activity on a specific region of interest by learning from EEG signal recorded simultaneously, inside the fMRI scanner. Indeed, the method proposed in (Meir-Hasson et al., 2014), uses a ridge regression model with a *ℓ*_2_ regularisation, based on a time/frequency/delay representation of the EEG signal from a single channel. Results show a good estimation of the BOLD signal in the region of interest, but the use of the fMRI neurofeedback in this study is only to serve the paradigm. The method aims at better targeting the amygdala in NF-EEG sessions.

Our challenge here, is to learn EEG activation patterns (see section 2.2) from hybrid (or bi-modal) NF-EEG-fMRI sessions (Perronnet et al., 2017), and improve the correlation with NF-EEG-fMRI of NF scores using EEG signal only. The motivation of this is multiple: since we are considering a new kind of data, we want to provide a simple method characterising NF-EEG-fMRI in EEG, leading to understandable model to confirm existing relations between EEG and fMRI in neurofeedback scores, or to discover new relationships. Neurofeedback features in fMRI come from the BOLD activation in one or more region of interest. We propose an original alternative to source reconstruction in the context of neurofeedback by taking advantage of the recent and unique dataset of NF-EEG-fMRI we have access to. Indeed we directly intent to predict NF scores, without dealing with source reconstruction or spatial filtering to estimate BOLD-fMRI signal first on a specific region of interest, as proposed by a previous approach (Noorzadeh et al., 2017). To our knowledge, this problem of prediction of hybrid neurofeedback scores (without source reconstruction) is new, and has not yet been explored in the literature. Also we want the activation pattern to be applicable in real-time when using new EEG data. The main objective of this paper (Figure 1) is to design a method able to exploit EEG only, and predict an NF score of quality comparable to the NF score that could be achieved with a simultaneous NF-EEG-fMRI session. The approach is based on a machine learning mechanism. During a training phase, both EEG and fMRI are simultaneously acquired and used to compute and synchronise, in real time, NF-EEG and NF-fMRI scores, both being combined into an hybrid NF-EEG-fMRI score (Mano et al., 2017). EEG signals and NF scores are used to learn activation patterns. During the testing phase, the learned NF-predictor (also called activation pattern) is applied to unseen EEG data, providing simulated NF-EEG-fMRI scores in real time. Sparse regularisation is exploited to build a model called NF-predictor. The model used for the NF-predictor uses an adapted prior for brain activation patterns, using a mixed norm giving a structured sparsity, to spatially select electrodes and then select the most relevant frequency bands.

**Figure 1.**
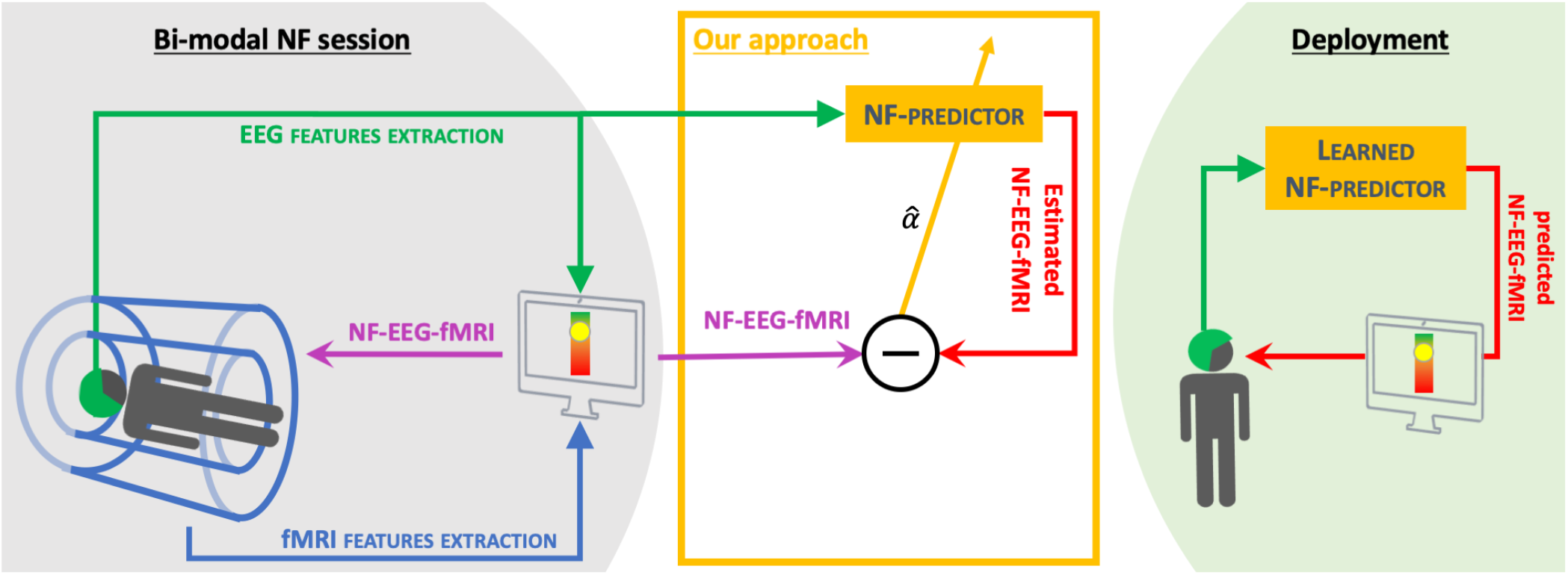
Objective: From bi-modal neurofeedback sessions (NF-EEG-fMRI) (see (Perronnet et al., 2017) or section 3 for more details), we propose a method to learn a NF-predictor. The final goal of this method is to propose NF sessions using EEG only, with the quality of bi-modal NF sessions. Therefore reducing the use of fMRI.

In section 2 we present the proposed model and the methods used to solve it. Then we will experiment our learning model on neurofeedback sessions with motor imagery task, which unique data are presented in section 3. Section 4 presents results on a dataset of 17 healthy subjects with 3 NF sessions of motor imagery each, one is used to learn the model, and the two others are used to test the model. Section 5 provides a discussion of the proposed framework.

## 2 PROBLEM AND METHOD

Considering that, during a learning phase, we have access to reference scores *y*(*t*) and a temporal representation (potentially non-linear) of EEG signals ***X*** (called a design matrix, presented in section 2.1); the approach consists in choosing a vector of parameters ***α*** such that *y*(*t*) ≈ *q*(***X*** (*t*), ***α***) for all *t*, where *q* is some parametric function. ***α*** is a matrix matching the size of ***X*** (*t*), here we consider

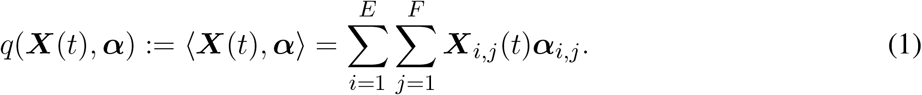

To determine this parameter ***α***, a regularisation is used to select an optimal parameter 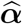 that fits the training data (see Figure 2 and section 2.3), while avoiding over-fitting, as detailed in Section 2.2.

**Figure 2.**
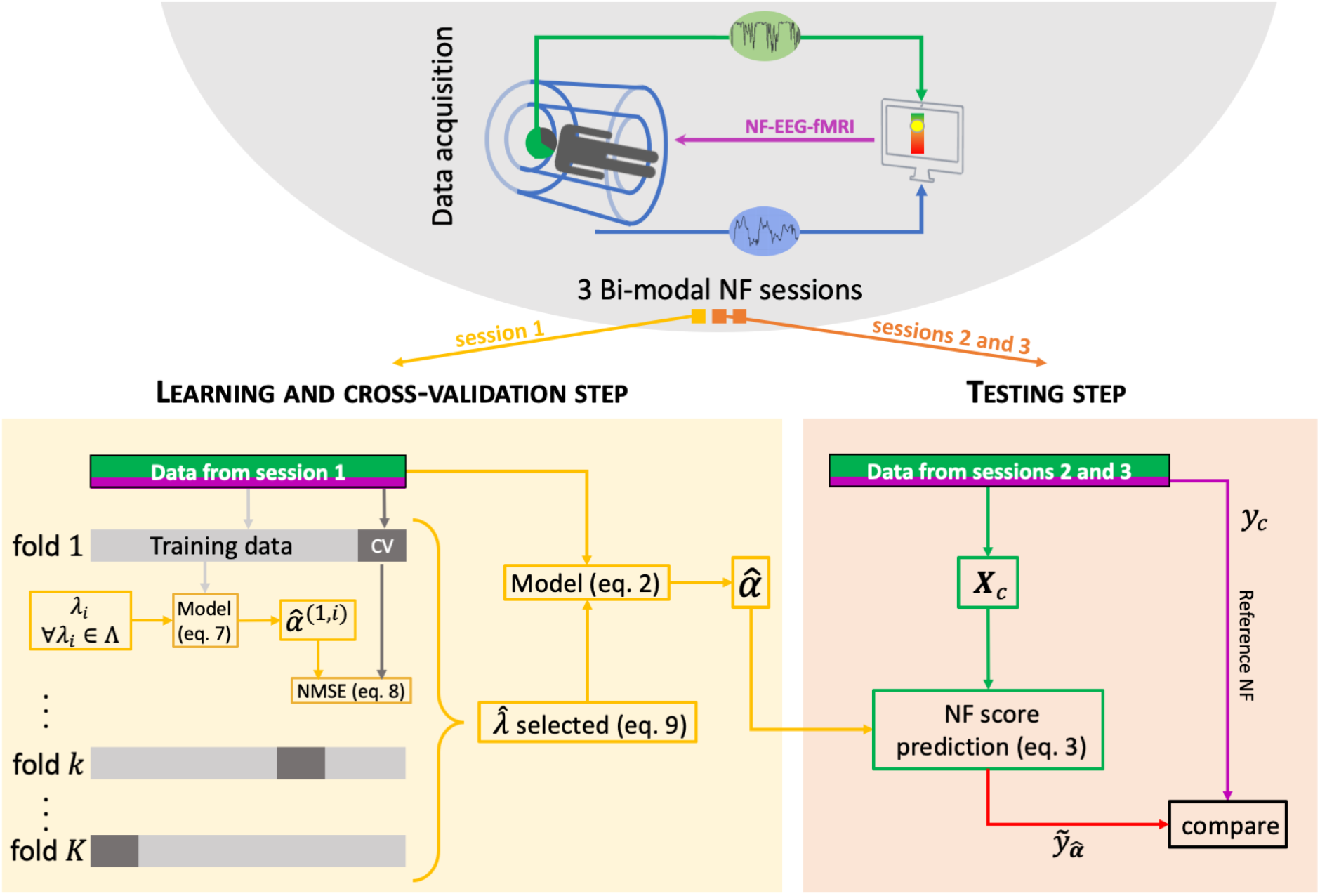
Machine learning scheme. For each subject, a bimodal neurofeedback session (NF-EEG-fMRI session 1 here) is used for the learning step, then the learned activation pattern 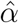 is apply to the other sessions (2 and 3) for the testing step. The learning data are split *K* times into a training set (90% of the learning set) and a cross-validation (CV) set (10% of the learning set). The optimal 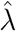 parameter is the one minimising the variance and the bias in the learning step.

Only a few brain regions are expected to be activated by a given cognitive task, therefore the electrodes configuration is said to be spatially sparse. However frequency bands of each electrodes are not necessarily sparse, and might even be smooth depending on the frequency band sampling.

From here, we will use the following notations:

- *y*_e_(*t*) ∈ ℝ, ∀*t* ∈ {1, …,*T*} are the *T* neurofeedback scores estimated from EEG signals (noted S_EEG_ ∈ ℝ^*E*×*T*_EEG_^), measured from *E* electrodes during *T*_EEG_ samples of time.
- *y*_f_(*t*) ∈ ℝ, ∀*t* ∈ {1,…,*T*} are the *T* neurofeedback scores extracted from Blood Oxygen Level Dependent imaging (BOLD) signal of functional-MRI acquisitions S_fMRI_ ∈ ℝ^*V*×*T*_fMRI_^, with *V* the number of voxels and *T*_fMRI_ the number of acquired volumes.
- *y*_c_(*t*) = *y*_e_(*t*) + *y*_f_(*t*) ∈ ℝ, ∀*t* ∈ {1,…, *T*} a combination of both NF scores (more details are provided in section 3).
- *y*(*t*) ∈ ℝ, ∀*t* ∈ {1, …, *T*} is a set of neurofeedback scores that can be *y*_e_, *y*_f_ or *y*_c_.

First, to build our predictor, relevant information from EEG data need to be extracted and organised to form what we call a design matrix.

### 2.1 Structured design matrices from EEG signal

The design matrix ***X***_0_ ∈ ℝ^*T*×*E*×*B*^, where *E* is the number of electrodes and *B* the number of frequency bands, contains relevant information extracted from the EEG signal. Each temporal matrix of ***X***_0_, ***X***_0_(*t*) ∈ ℝ^*E*×*B*^ ‘∀*t* ∈ {1;…; *T*} is a frequency decomposition corresponding to the past 2 seconds of *S*_EEG_. We used a Hamming time window of 2 seconds, to estimate the average *power* of each frequency band *b* ∈ {1;…; *B*} (defined below) on each channel ∈ {1;…; *E*}. Each time window of EEG signal is overlapped by 1.75 seconds (0.25 seconds shift), to match with the 4Hz sample of the ***y*** values. The B frequency bands have an overlap of 1 Hz with the next band, and are defined between a minimum frequency *b_min_* Hz and a maximum frequency *b_max_* Hz (see section 3). We chose to use several relatively narrow frequency bands to let the model select the relevant bands for each electrodes. Furthermore it has been suggested (de Munck et al., 2009; Rosa et al., 2010) to use different frequency bands when working with coupling EEG-fMRI data.

The model also has to be able to predict *y*_f_ scores, derived from BOLD signal (see section 3). There is no linear relationship between BOLD signal and average power on frequency bands from EEG signal. Therefore, to better match *y*_f_ scores, we decided to apply a non-linear function to ***X***_0_, used in fMRI to model BOLD signals (Pedregosa et al., 2013; Lindquist et al., 2009), the canonical Hemodynamic Response function (HRF). We convolved ***X***_0_ on its temporal dimension with the HRF, formed by 2 gamma functions, for a given delay of the first gamma function to compensate the response time of BOLD signal, as suggested in (Meir-Hasson et al., 2014; Moosmann et al., 2008). The HRF will temporally smooth and give a BOLD-like shape to the design matrix and increase a potential linear relationship between *y*_f_ and design matrix. Since HRF is known to vary considerably across brain regions and subjects (Handwerker et al., 2004), it is therefore recommended to consider different delays, but also to chose a range of values corresponding to the task asked. For the type of task addressed in the experimental part, the observed delay is around 4 seconds, therefore we convolved X0 with 3 different HRFs leading to 3 new design matrices ***X***_3_, ***X***_4_, ***X***_5_ with respectively peak locations of 3, 4 and 5 seconds. We started from the canonical HRF and changed the peak parameter (respectively using 3 4 and 5) to induce the 3 different HRF used to induce time delay to the initial design matrix ***X***_0_. By doing so, we keep the total of design matrices of a reasonable size

Those design matrices are concatenated in their 2nd dimension to form the ***X***_*c*_ ∈ ℝ^*T* ×*M* ×*B*^ matrix, with *M* = 4 * *E*. Therefore, for each time *t*, ***X***_*c*_(*t*) = [***X***_0_(*t*); ***X***_3_(*t*); ***X***_4_(*t*); ***X***_5_(*t*)]. We also denote ***X***_*d*_(*t*) = [***X***_3_(*t*); ***X***_4_(*t*); ***X***_5_(*t*)] the design matrix of the different delays.

### 2.2 Optimisation

EEG data are now represented into a structured design matrix ***X***_*c*_, we can search for a weight matrix 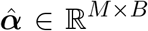, such that 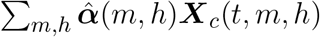 estimates as well as possible the NF score *y*(*t*), ∀*t* ∈ {1;…; *T*}. Note: the methodology is presented for the design matrix ***X***_*c*_, but can be used for ***X***_0_ or ***X***_*d*_.

To identify the 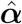, called activation pattern, we propose the following strategy, which consists in learning, for a given subject and a NF session, the optimal 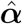 by solving the following problem:

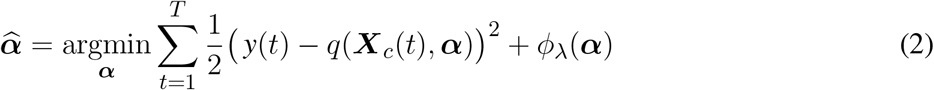

with *φ*_λ_ a regularisation term, λ a weighting parameter for the regularisation term. This 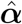 is then applied to a design matrix 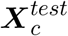 from a new session, to predict its NF scores

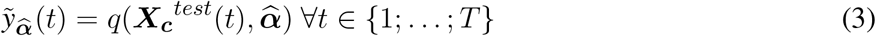

Equation (2) is of the form argmin(*g*_1_ (***α***)+*g*_2_ (***α***)) and its resolution can be done using the Fast Iterative Shrinkage Thresholding Algorithm (FISTA) (Beck and Teboulle, 2009), which is a two-step approach of the Forward-Backward algorithm (Combettes and Wajs, 2005) making it faster. FISTA requires the same conditions as the Forward-Backward algorithm, meaning a convex differentiable with Lipschitz gradient term *g*_1_ and a convex term *g*_2_ that is not necessarily differentiable but smooth enough to make its proximal map computable.

Here 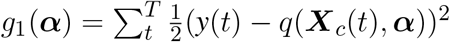 is a sum of convex and differentiable functions with

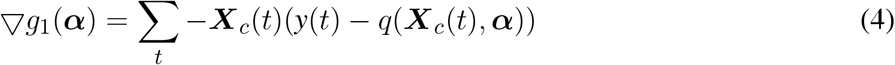

since 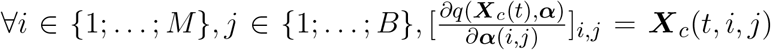. By representing ***X***_*c*_(*t*) and ***α*** as vectors of size *M * B*, we can easily note that 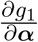 is a sum of Lipschitz functions. Therefore, the Lipschitz constant of 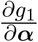 is 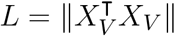 with *X_V_* ∈ ℝ^*T*M*B*^ the vectorised version of ***X***_*c*_.

The NF-predictor uses structured design matrix to have a better control on the interpretation of results and to better optimise the weights 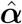. Therefore we have to adopt an optimisation strategy coherent with this structure. The activation pattern of the NF-predictor:

1. has to be spatially sparse to regulate the model as EEG signals are noisy and to select the most relevant electrodes on each frequency bands,
2. has to be smooth across different overlapped frequency bands,
3. has to allow non-relevant frequency bands to be null.

The term *g*_2_ is the prior term. Here, for *g*_2_(***α***) = *φ*_λ_(***α***), we chose to use a *ℓ*_21_ mixed norm (Ou et al., 2009) followed by a *ℓ*_1_-norm (noted *ℓ*_21+1_-norm in (Gramfort et al., 2011)) to fit all structure conditions mentioned above.

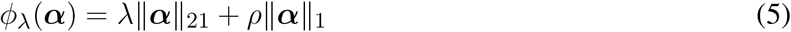

with *ρ* ∈ ℝ^+^ and λ ∈ ℝ^+^. We chose not to estimate the parameter *ρ*, to keep computation time reasonable. Indeed, *ρ* weights the induced spatial sparsity over EEG channels, and we chose to fix this parameter for all subjects, as we hypothesis that there is no reason, for the number of electrodes involved in the activation pattern, to significantly change between subjects. However the estimation of λ parameter is needed (since we do not have hypothesis on its behaviour) and presented in the next section. The *ℓ*_21_ mixed norm that writes 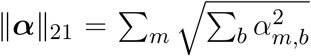 satisfies conditions 1) and 2). The *ℓ*_1_ norm defined as ||***α***||_1_ = Σ_*m,b*_ |*α_m,b_*| satisfies condition 3) since *ℓ_p_* normswith *p* ≤ 1 are known to promote sparsity. The last key point of FISTA algorithm is the proximal map associated to the *ℓ*_21+1_ norm Prox_*ℓ*21+1_: ℝ^*M×B*^ → ℝ^*M×B*^, *β* ↦ argmin_*α*_ (*ϕ*_λ_ (***α***) +1/2||*β* – *α*||^2^), defined as

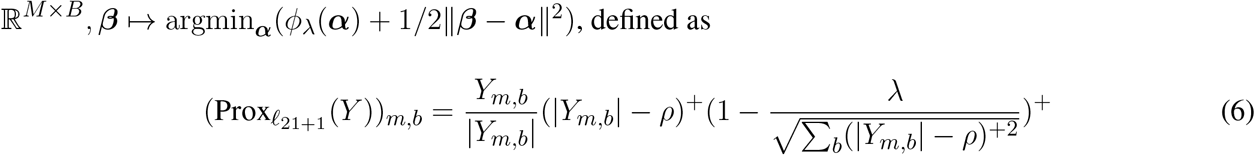

with operator (.)^+^ = *max*(., 0). One can note that by cancelling either the λ parameter or the *ρ* parameter, we retrieve the proximal map associated to the *ℓ*_21_ (when *ρ* = 0) and to the *ℓ*_1_ norm (when λ = 0), which demonstrations can be found in the appendix of (Gramfort et al., 2012). For the stopping criterion of FISTA, a large enough number of iteration has been used, allowing the model to converge before reaching the last iteration. All elements and conditions are gathered to run the FISTA algorithm.

### 2.3 λ parameter selection

The parameter λ is important in the optimisation problem and we decided to estimate it automatically. The following process chooses the best λ among a list of Λ = {λ_1_;…; λ_*l*_} sorted in increasing order. First of all, the data must be split into 2 sets. In our case subjects have 3 NF-EEG-fMRI sessions: one session is used as the learning set, and another NF-EEG-fMRI session is used as the testing set (see Figure 2). For each value λ_*i*_ of Λ, the learning set, formed by *T* neurofeedback scores with their associated design matrices, is divided *K* = 50 times into a training set of indices *R_k_*, representing 90% of the *T* data, and a cross-validation set *CV_k_* composed by composed by the remaining 10% of the learning set. A model 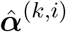 is estimated on the training dataset *k* composed by *R_k_* neurofeedback scores *y*(*j*) and the associated design matrices ***X***_*c*_(*j*) with λ_*i*_, i.e.:

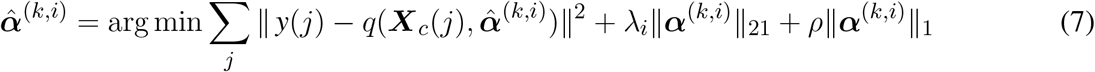

with *j* ∈ *R_k_* and ***X***_*c*_(*j*) ∈ ℝ^*M×B*^. For the current λ_*i*_ evaluation, we stop the process when 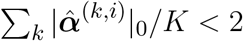. There is no need to investigate the next λ_*i*_, the current one is sparse enough, and the next one might lead to null models.

We then apply the model 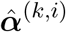 to the corresponding cross-validation set of *CV_k_* NF scores, to obtain estimated values of 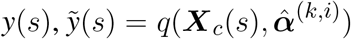, with *s* ∈ *CV_k_*. For each one of the 50 partitioning into training and cross-validation sets, we computed the normalised mean squared error NMSE for a given set of data {*y*, ***X***_*c*_}, for the training sets and the cross-validation sets.

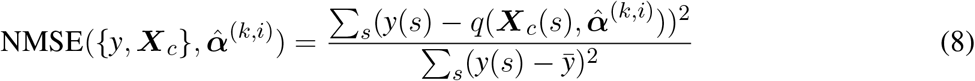

with 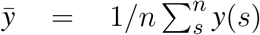. The optimal 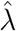 parameter is defined as the one minimising the error during training and cross-validation. Considering only the errors from the training set NMSE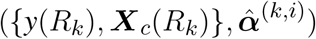 would introduce bias, and considering only the error of the crossvalidation set NMSE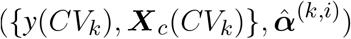, would introduce variance. Then the optimal 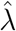 is the λ_*i*_ that minimises:

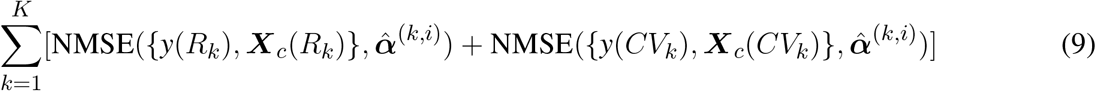

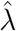 parameter is the optimal parameter used for the model estimation. If there are several candidates, to favor sparsity, the larger of these candidate is chosen.

## 3 DATA ACQUISITION AND PRE-PROCESSING

We used an existing dataset presented in (Perronnet et al., 2018), composed of 17 healthy subjects that were scanned using the hybrid Neurofeedback platform from Neurinfo (Rennes, France) coupling EEG and fMRI signal (Mano et al., 2017). Data are now available online in BIDs format on openneuro: https://openneuro.org/datasets/ds002338 (Lioi et al., 2019). A 64-channel MR-compatible EEG solution from Brain Products (Brain Products GmbH, Gilching, Germany) has been used, the signal was sampled at 5kHz, FCz is the reference electrode and AFz the ground electrode. For the fMRI scanner, we used a 3T Verio Siemens scanner with a 12 channels head coil (repetition time (TR) / echo time (TE) = 2000/23ms, FOV = 210 × 210mm^2^, voxel size = 2 × 2 × 4mm^3^, matrix size =105 × 105 with 16 slices, flip angle = 90°). All subjects are healthy volunteers, right-handed and had never done any neurofeedback experiment before. They all gave written informed consent in accordance with the Declaration of Helsinki, as specified in the study presenting the data (Perronnet et al., 2017). They all had 3 NF motor imagery sessions of 320 seconds each, after a session dedicated to the calibration. One session consists in 8 blocks alternating between 20 seconds of rest, eyes open, and 20 seconds of motor imagery of their right hand. The neurofeedback display was uni-dimensional (1D) for 9 subjects (Figure 3 left), and bi-dimensional (2D) for 8 subjects (Figure 3 middle). For both, the goal was to bring the ball into the dark blue area (Perronnet et al., 2018).

**Figure 3.**
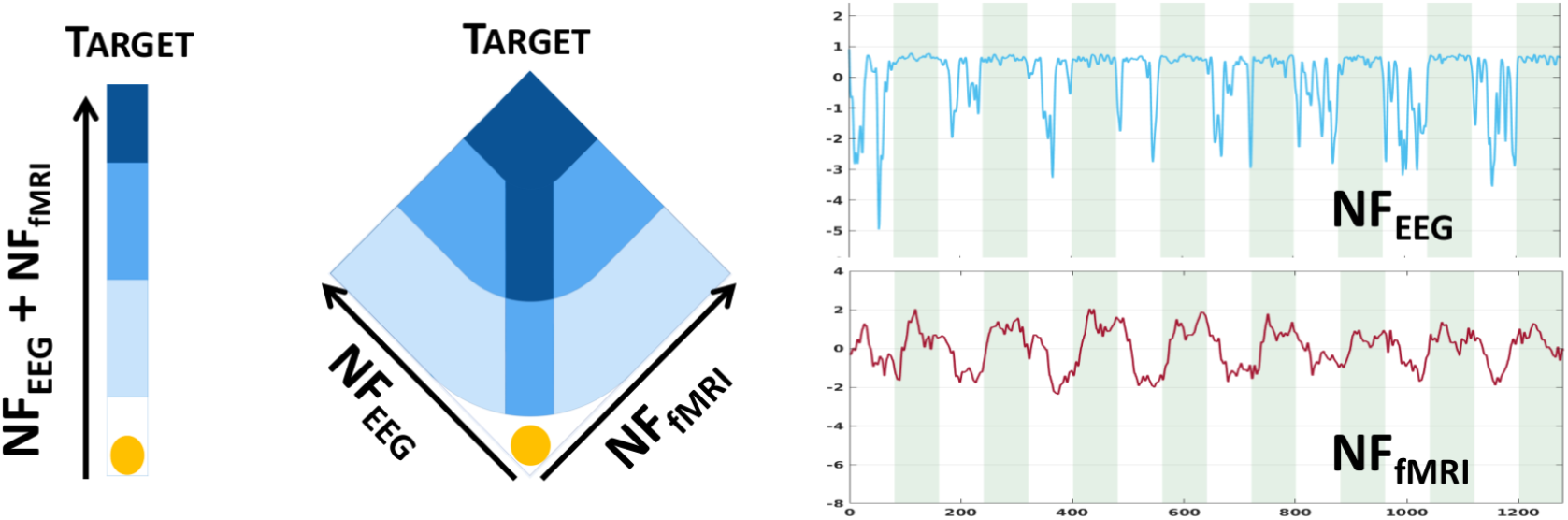
Bi-modal neurofeedback metaphors (1D on the left, 2D on the middle), displayed during sessions (Perronnet et al., 2018). 1D: the ball’s position represents the sum of *y*_e_ and *y*_f_. 2D: the left axis represents the *y*_e_ and the right axis represents the *y*_f_ scores. The 2 plots on the right show NF scores from EEG and from fMRI, green areas are task, white areas are rest. The goal is to bring the ball in the dark blue area.

NF scores *y*_e_ and *y*_f_ being from different modalities, were standardised before summing to form *y*_c_. In this study NF scores refer to standardised scores, except when they are predicted. For this study, *y*_e_ have been computed from the commonly used, in neurofeedback, the Laplacian operator, centred around the region of interest, channel C3 here. For each time interval *I_t_* the spatial filtering is noted Lap(C3, *I_t_*). The temporal segments *I_t_* are spaced by 250ms, and a length of 2 seconds (therefore an overlapping of 1,75 seconds), as for the design matrix construction. The power of the frequency band [8Hz - 30Hz] is then extracted via the Power Spectral Density function PSD:

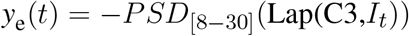

One may note the presence of the minus operator used here, for the sake of coherence with *y*_f_ (Figure 3 right).

The neurofeedback scores *y*_f_ have been computed from the maximal intensity of BOLD signal covering the right-hand motor area and the supplementary motor area, one score is computed per volume acquired (i.e 1 per second). Then scores *y*_f_ are re-sampled and smoothed (using a Savitzky-Golay filter, known to avoid signal distortion) to fit the 4Hz *y*_e_ scores (*T* = 1280).

An active set have been selected on design matrices to avoid potentially correlated noise, due to head movement during resting blocks, obstructing signal from channels of interest. Indeed in coupling EEG-fMRI acquisitions, subjects are lying into the MRI scanner, therefore outer electrodes can be in contact with the bed or holds. We excluded outer electrodes and kept 28 electrodes, the 3 central lines have 7 electrodes (FCz is the reference), 3 frontal electrodes and 3 posterior electrodes.

Potential outliers in the design matrices (i.e. observations > mean±3std) were thresholded in the NF-EEG-fMRI session used as learning set, and bad observations from annotations on the EEG signal were removed as their corresponding NF scores. For frequency dimension of the design matrix ***X***_0_ construction (cf section 2), and therefore for the other design matrices, we chose *b_min_* = 8, *b_max_* = 30 to cover alpha and beta frequency bands involved in motor tasks. We considered *B* =10 frequency bands, leading to bands of 3 Hz wide with an overlap of 1Hz.

As mentioned in the previous section, for the regularisation of the NF-predictor, we used 15 values of λ_*i*_ from 100 to 3000 and fixed the parameter *ρ* = 1500.

## 4 EXPERIMENTS AND RESULTS

As said in the previous section, we have 3 neurofeedback sessions per subjects. For each subjects, we will consider 1 session as learning set, and the 2 others as testing sets. Leading to 3 different learning sets, and 6 different testing sets per subjects.

### 4.1 Experiments and validation

We tested different NF-predictors for the prediction of different NF scores, ∀*t* ∈ {1;…; *T*}:

1. 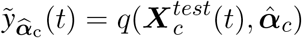 with 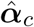 (eq. 2), learned from ***X***_c_ and *y*_c_
2. 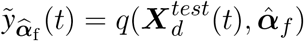 with ***X***_d_ = [***X***_3_; ***X***_4_; ***X***_5_], and 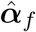 (eq. 2), learned from ***X***_*d*_ and *y*_f_
3. 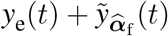 using NF-predictor 2

The advantage of the last NF-predictor is the possibility of using the 2D score visualisation (Figure 3) to display NF scores. Also in this case, only NF-fMRI scores has to be learned. We run the following experiments to test our different NF-predictors:

- Model validation: we used the learning set to assess if the different NF-predictors could model accurately the NF scores. For each subject and each NF-predictor, we estimated correlations with the reference NF score to quantify the quality of prediction. As a reference, we compared to the correlation of *y*_e_ to *y*_c_, which is part of the *y*_c_ score, to assess if, in the validation process, the model can better predict *y*_c_ scores than *y*_e_.
- Model prediction: we apply the learned activation patterns to a new NF-EEG-fMRI recorded session (testing set, Figure 2). For each subject and each pair of session (6 learning/testing pair per subject), we compared correlations between NF-predictors 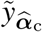 and 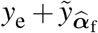 to reference score *y*_c_, and between the prediction of 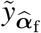 to *y*_f_.
- To observe the captured structure in the activation pattern 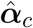 and 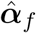 respectively learned to predict *y*_c_ and *y*_f_, we re-shaped the averaged activation patterns of a subject over sessions, into the matrices corresponding to the design matrices defining ***X***_*c*_ respectively ***X***_*d*_ (see section 2.1), and displayed results of the first dimension (electrodes) and of the second dimension (frequency bands).

### 4.2 Results

#### 4.2.1 Model and prediction

The model validation of the NF-predictors (i.e. the learned activation patterns 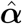 are applied to the learning set) results are shown at Figure 4 left side. Pearson’s correlation coefficients between the prediction and the ground truth are computed for each of the 3 learning sessions and for all subjects. Correlations are very high (> 0.8) for the 2 NF-predictors of *y*_c_ scores, letting think that the model is adapted to the problem of NF prediction. Indeed, the model could fit NF-fMRI scores using only EEG signals information with a median correlation of r = 0.81 with *y*_f_.

**Figure 4.**
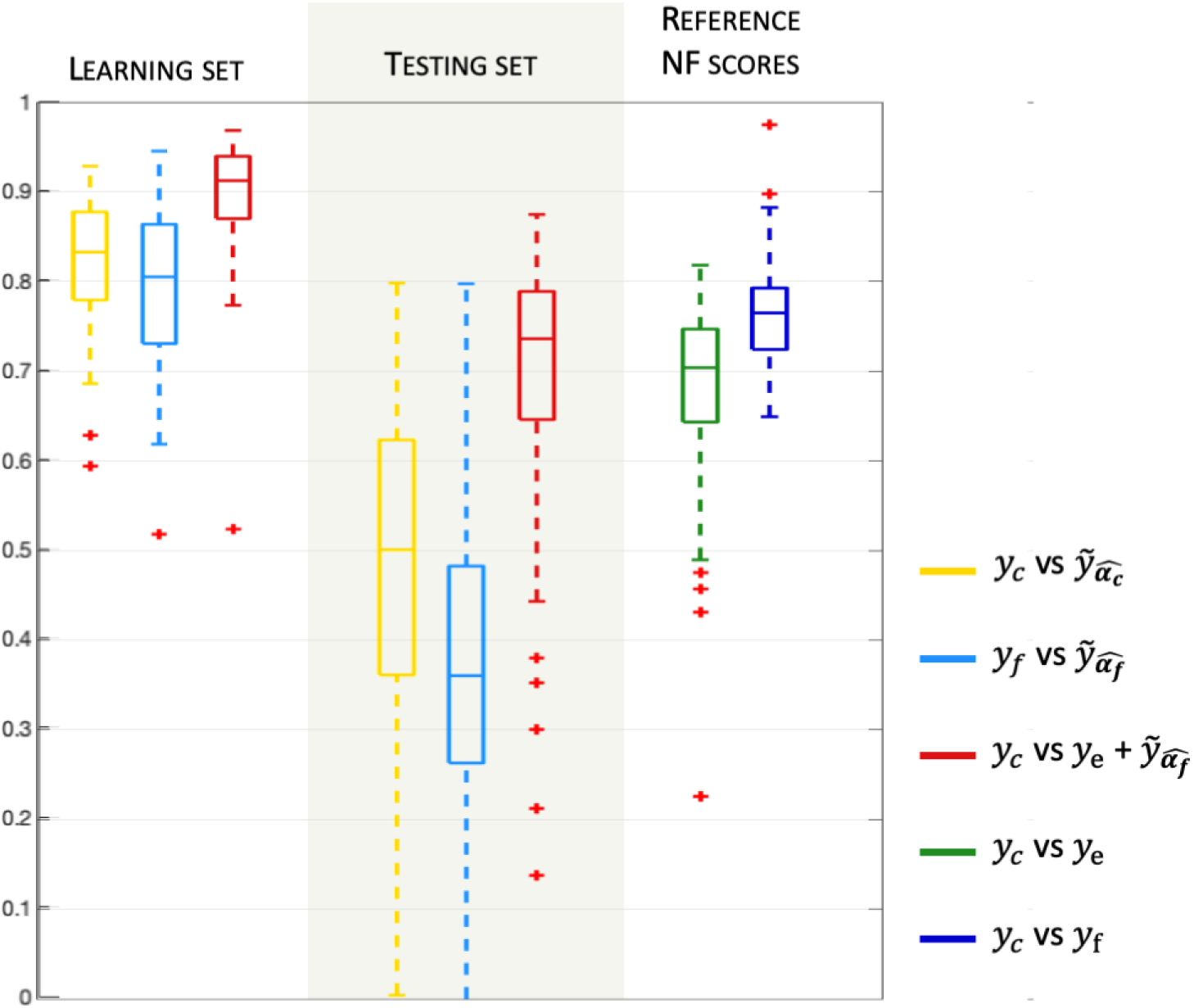
Model validation and prediction. Boxplots (median and quartiles) of Pearson’s correlation coefficients over all subjects and sessions, between NF-predictors and *y*_c_ or *y*_f_. The right part of the plot indicates the correlation of the reference NF scores *y*_e_ and *y*_f_ versus the corresponding reference bi-modal NF scores *y*_c_(= *y*_e_ + *y*_f_).

For the evaluation of the model prediction, the learned activation patterns are applied to the testing sets i.e. the 2 other unseen NF sessions, for each one of the 3 NF sessions. The significance of the differences between the correlations to the reference NF scores is assessed by applying a Fisher transformation to the correlation coefficients. The p-values obtained with Student’s t-test, after this transformation, indicate the significance of the differences between the correlation coefficients distributions. Table 1, shows that 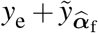, whose median correlation is r = 0.74, better predicts *y*_c_ than *y*_e_ only, despite the fact that *y*_e_ is part of *y*_c_. The paired t-test, gives a p-value *p* = 6.6e-4 (*t* = 3.52), meaning that the prediction of NF-fMRI scores by 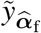 significantly adds information to *y*_e_. Predicting *y*_c_ scores with 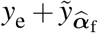 is also better than 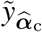 (*p* = 2e-42, *t* = 23.17), which is expected since the models predicts NF-fMRI scores and directly use NF-EEG scores (with all the potential noise coming from the EEG measures) which are part of the reference score, as *y*_e_ is in 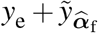, and therefore correlates well with *y*_c_. Furthermore the proposed model is able to predict *y*_f_ scores with a fair correlation of 0.36 in median and 0.35 in average. Thus predicting NF-fMRI scores, instead of predicting bi-modal NF scores, seem to be the best way of predicting those bi-modal NF scores. Figure 5 gives examples of predictions of *y*_c_ with 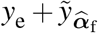 and 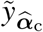.

**Figure 5.**
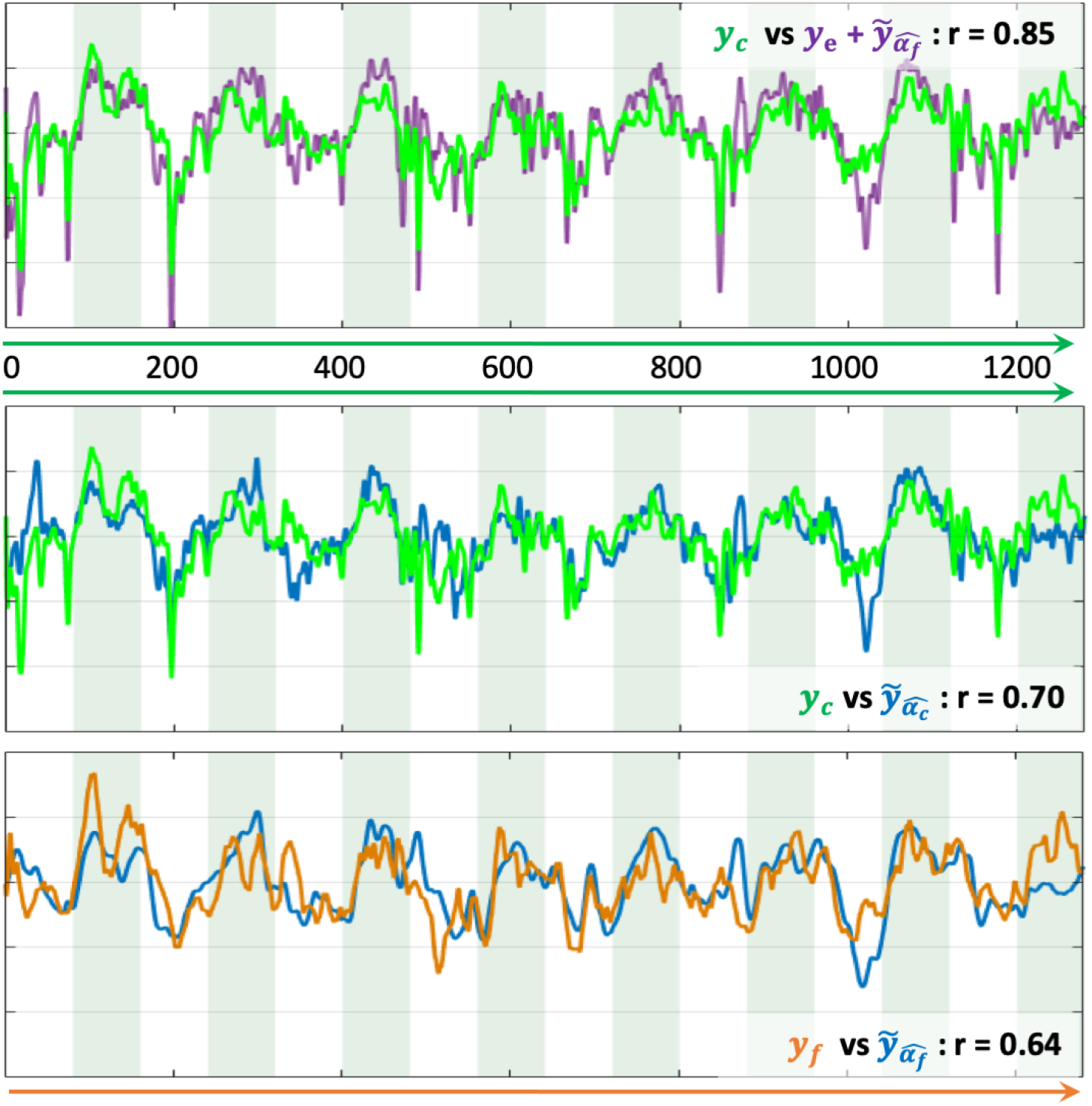
Examples of prediction of NF scores. The x-axis is the temporal axis in milliseconds. Vertical bands indicate the rest and the task blocks. The correlation coefficient *r* indicates the correlation between each pair of time-series NF scores. Top: prediction of *y*_c_ with 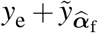. Middle: prediction of *y*_c_ with 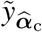. Bottom: prediction of *y*_f_ with 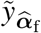.

**Table 1.** Model prediction: Pearson’s correlation coefficients over all subjects and sessions, between NF-predictors and *y*_c_. The table shows p-values of t-test, the hypothesis is: models in columns correlates better with *y*_c_ than models in rows.

| Correlation ($y_c, \cdot$) | paired t-test | | |
| --- | --- | --- | --- |
| | $\tilde{y}_{\hat{\alpha}_c}$ | $y_e + \tilde{y}_{\hat{\alpha}_f}$ | $y_e$ |
| $\tilde{y}_{\hat{\alpha}_c}$ | N.A. | $t = -23.17$<br>$p = 1$ | $t = -12.76$<br>$p = 1$ |
| $y_e + \tilde{y}_{\hat{\alpha}_f}$ | $t = 23.17$<br>$p = 2.e-42$ | N.A. | $t = 3.52$<br>$p = 6.6e-4$ |
| $y_e$ | $t = 12.76$<br>$p = 9e-23$ | $t = -3.51$<br>$p = 1$ | N.A. |

It also illustrates that, even if the correlations of 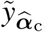 are lower than 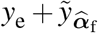 (Table 1), 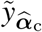 can predict correctly the reference score *y*_c_.

During the learning session (i.e. the NF session used to learn the predictor), subjects might unevenly focus on their fMRI or EEG feedbacks, and when subjects focus more on NF-fMRI than on NF-EEG, the EEG-signals might lose coherence with respect to the NF-fMRI scores. Even though, the EEG signals could predict NF-fMRI scores with a correlation of 0.36 in median and mean 0.35 (cf Figure 4, which is a fair correlation between such different modalities. Examples of NF prediction are given at the bottom of Figure 5, the plot shows the prediction 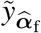 of NF-fMRI scores on a testing set.

#### 4.2.2 Activation patterns

We now focus on the learned models to evaluate the sparsity, over sessions and subjects and observe the dispersion of the learned patterns over sessions and frequency bands of a subject. For each subjects and each learning session, we computed the proportion of zero coefficients of the activation patterns (Table 2). By construction of the design matrix 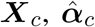 and 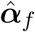 can be split into different activation patterns, as shown in columns of Table 2. The models could select relevant coefficients of the design matrices to predict reference NF scores. In average, there is 87 non-zeros coefficient on 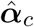 to predict *y*_c_, and 57 non-zeros coefficients on 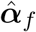 to predict *y*_f_.

**Table 2.** Sparsity of the learned models 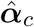 and 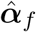. Proportion of zero coefficients in the learned models.

| | $\hat{\alpha}_0$ | $\hat{\alpha}_3$ | $\hat{\alpha}_4$ | $\hat{\alpha}_5$ | all |
| --- | --- | --- | --- | --- | --- |
| $\hat{\alpha}_f$ | x | $0.92 \pm 0.03$ | $0.96 \pm 0.03$ | $0.91 \pm 0.03$ | $0.93 \pm 0.04$ |
| $\hat{\alpha}_c$ | $0.87 \pm 0.05$ | $0.92 \pm 0.03$ | $0.96 \pm 0.02$ | $0.93 \pm 0.03$ | $0.92 \pm 0.05$ |

To display the activation patterns of a subject over its 3 sessions, we denote 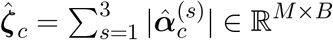 the absolute activation pattern of 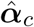 (respectively 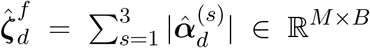 for 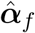), and 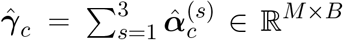 the average activation pattern (respectively 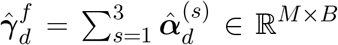). We can display heat maps for each of the 4 absolute activation patterns at each electrodes 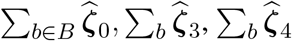 and 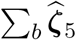 (Figure 6, top line of each panels); and colour maps of the 4 average patterns 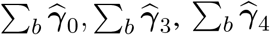, and 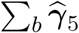 (Figure 6, bottom line of each panels) showing the sign of the strongest and most stable coefficients across all subjects and sessions. Activation patterns displayed at Figure 6, represent the dispersion of the learned parameters for one example subject over its 3 bi-modal NF sessions for which he received a bi-dimensional display. The subject used had a median correlation with the corresponding reference score of 0.44 for 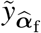, of 0.76 for 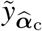 and of 0.81 for 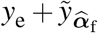. All maps present different distributions of the non-zero coefficients. As expected when learning *y*_c_ scores, the most intense heat map 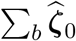 with a maximum value of 45, concentrates its non-zero values on C3 channel (above the righthand motor area) and corresponds to the design matrix ***X***_0_, directly extracted from EEG signal without non-linear temporal delay. The next heat map in intensity order is 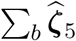 with two peaks with a parietal positioning above Pz and P4 channels, corresponding to the Brodmann area 7, involved in the visuo-motor coordination. As presented in an interesting study (Sitaram et al., 2017), this brain area is active during generalised neurofeedback when feedback is presented visually. In (Sitaram et al., 2017) it is also mentioned that this part of the cortex is part of the executive control network connected to the thalamus. The activation of the executive control network during motor imagery task is relevant, and indicates that the subject is trying to do the task. It is also interesting to observe that the main activation peaks of 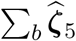 (Pz and P4) have opposite signs with the activation peak of 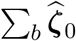 (above C3), suggesting a negative correlation between neural activation measured at C3 and the neural activation of posterior parietal Pz and P4 channels, part of the control network in neurofeedback. Panel B of Figure 6 presents comparable learned activation patterns to predict 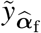, however more spread over sessions than for the model on panel A, even if the model as around 93% of null coefficients (Table 2). This suggests that, even for a same subject, the relation between EEG and fMRI changes over sessions, probably depending on the strategy used by the subject.

**Figure 6.**
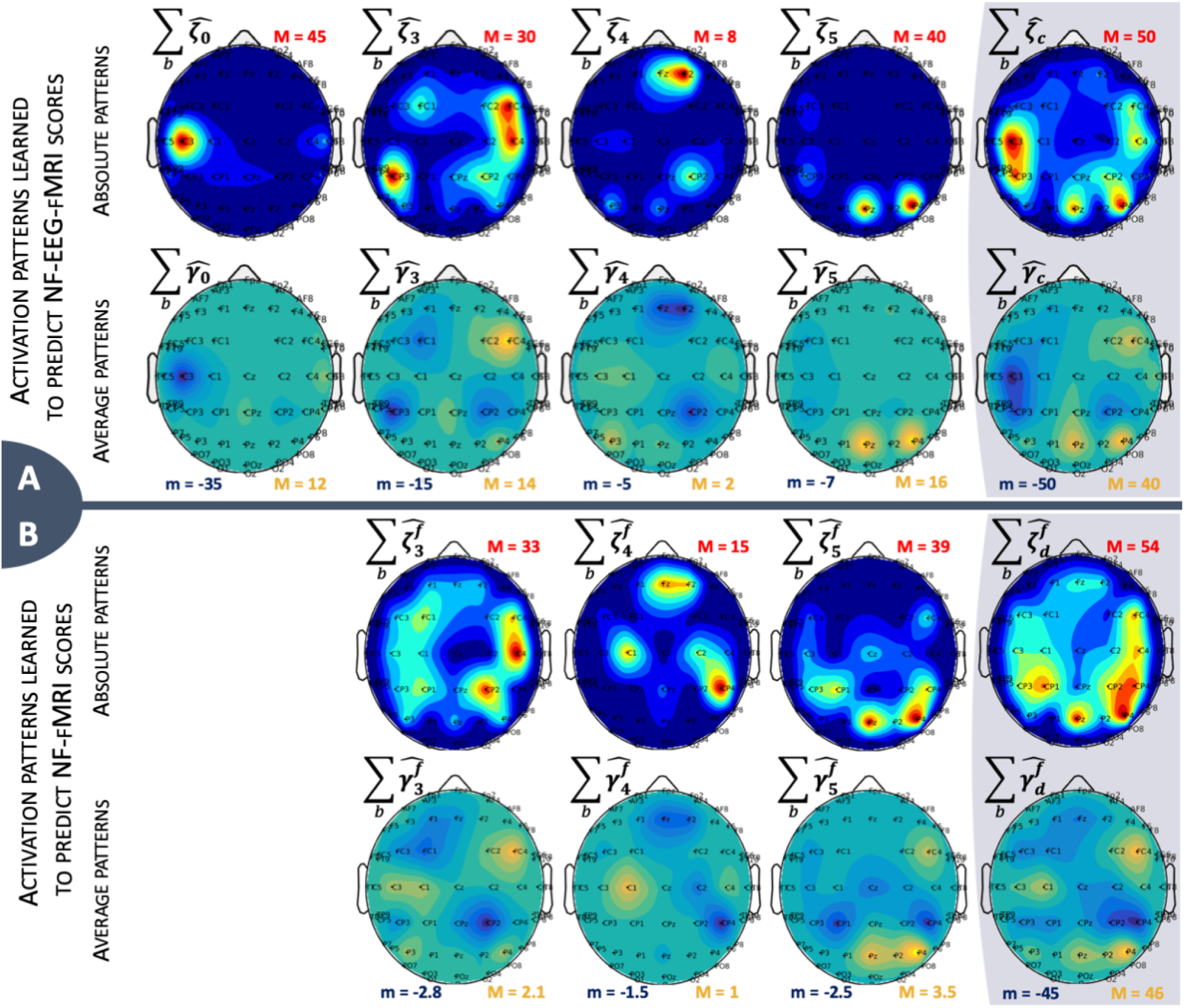
Activation patterns over sessions and frequency bands *b* for one example subject. There is two lines for each model, panel A represents activation patterns for 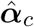 and panel B represents activation patterns for 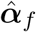. The last column is the sum of the other ones. Top line: Heat map representing the distribution of non-zero coefficients. Maximum value is indicated for each map with the red letter M, dark blue areas represent zero values. Bottom line: Average activation patterns, representing the sign of the main non-zero values across subject and sessions. Minimum and maximum values are indicated for each map with the letters m and M, green areas represent zero values.

At last, it is also possible to display the frequency profile of each average activation patterns. The 4 frequency profiles are 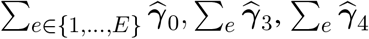 and 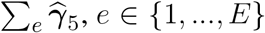, summing weights over electrodes. Figure 7 shows that the most used frequency bands over all sessions and electrodes are alpha band ([8-12] Hz) and lower beta band ([13-17] Hz), the last 4 frequency bands are not displayed as they only have null coefficients. We can observe the effect of *ℓ*_2_ regularisation which allows continuity in the frequency bands, and the effect of the subsequent *ℓ*_1_ regularisation which removed the smaller coefficients located in the high frequencies. When considering all activation patterns (Figure 7 right side), there is a change of sign between alpha and lower beta. Each activation pattern shows a different frequency profile, which, together with Figure 6, tends to demonstrate that patterns have complementary information.

**Figure 7.**
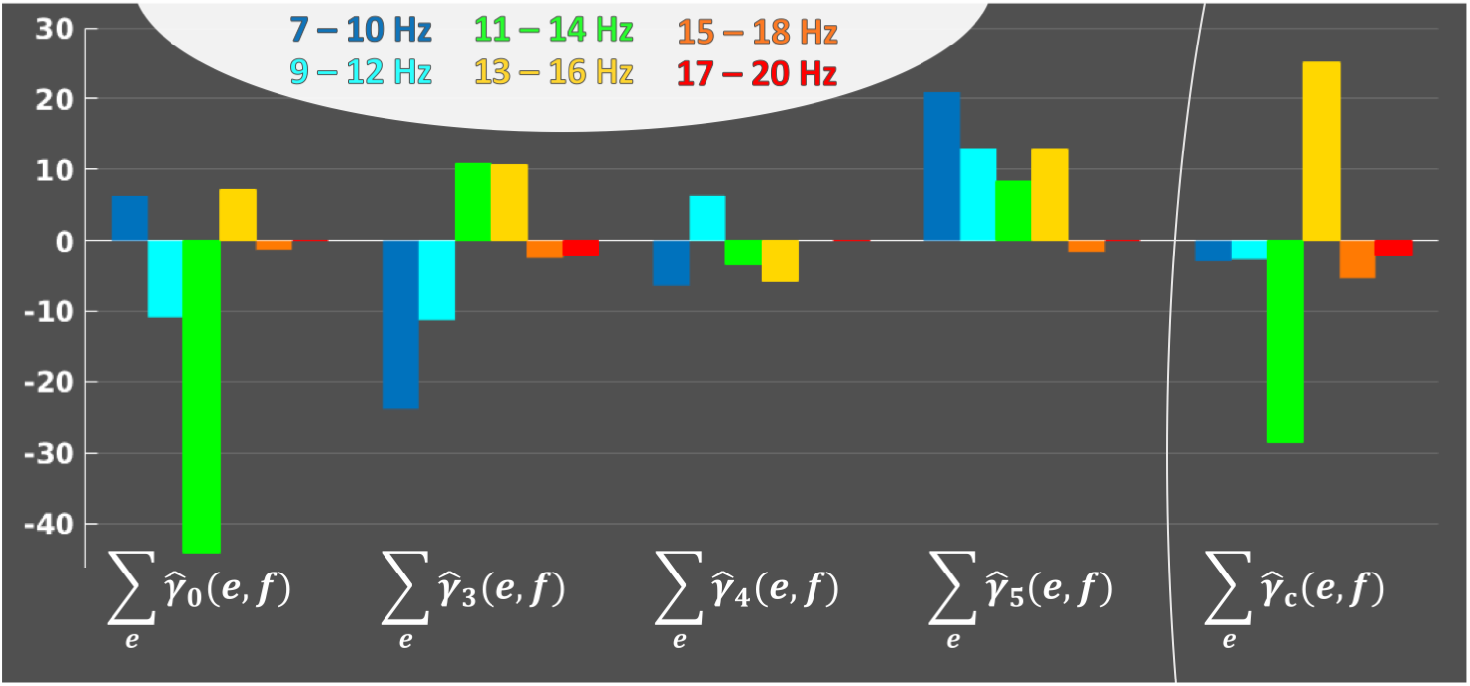
Frequency profiles, across subjects and sessions, of each average activation patterns, to represent the implication of each frequency bands in the activation patterns. *e* ∈ {1,…, *E*}. For each average activation pattern, the y-axis indicates the sum of weights over electrodes, for each frequency bands *f* ∈ {1,…, *B*} on the horizontal axis.

### 4.3 Ablation study

We run an ablation study to understand the impact of the non-linearity part of the model, obtained by the convolution of different HRFs inducing different delays for the prediction of NF-fMRI scores, by analysing results of the model without HRFs convolutions. Therefore, in this ablation study we used the exact same processes as described in section 4 (with a learning step and a testing step), except that we used the design matrix ***X***_0_ alone to learn the model parameters to predict *y*_c_ scores and *y*_f_ scores.

Results of the ablation study are displayed at Figure 8 and are to be compared with results of the proposed model presented at Figure 4. Figure 8 shows that during learning step, the prediction of 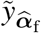, using *X*_0_ only, correlates with *y*_f_ with a correlation of only 0.48 in mean and median. This prediction is significantly improved by the use of the different HRFs functions (paired t-test, *t* = −14,64, *p* = 1*e*-26), as in the proposed model 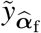 correlates with *y*_f_ with a correlation of 0.81 in median and mean. Therefore, it is natural to observe that the prediction of the bi-modal NF score by 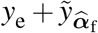 is also significantly improved by the use of non-linearity (paired t-test, *t* = −11,87, *p* = 4*e*-21). Figure 8 shows that for the testing step, the prediction of 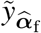, using *X*_0_ only, correlates with *y*_f_ with a correlation of 0.14 in mean and 0.15 in median, which is significantly lower than the proposed model (paired t-test, *t* = −9.56, *p* = 2*e*-18). The prediction of the bi-modal NF scores by 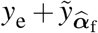 is also significantly lower than the proposed model (paired t-test, *t* = −2.74, *p* = 3*e*-3). Also, when using *X*_0_ only, the prediction of the bi-modal NF scores using 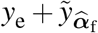 do not significantly improve over *y*_e_ alone (paired t-test, *t* = −0.69, *p* = 0.5).

**Figure 8.**
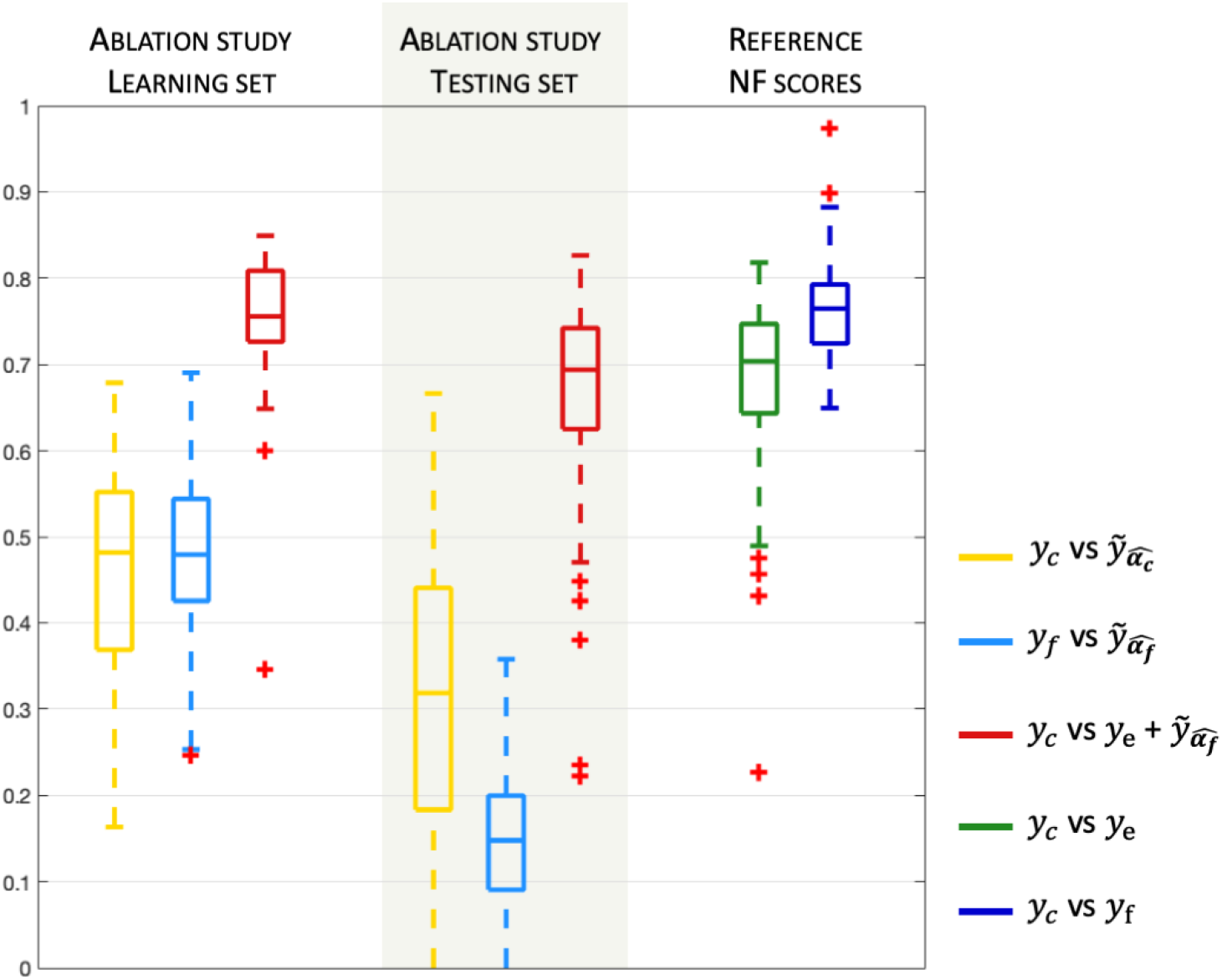
Ablation study. Removing non-linearity, keeping *X*_0_ only in the model to understand the importance of the non-linearity induced by the use of different HRF functions.

## 5 DISCUSSION AND CONCLUSION

The model validation supports that the optimisation strategy we chose for our problem is adapted to the model, as is the choice of the different design matrices. The ablation study supports the use of different non-linear delays to improve the prediction of NF-fMRI scores using EEG signal. The evaluation of the model prediction strongly suggests that predicting only NF-fMRI scores from EEG signal while applying a Laplacian on EEG signal appears to be the best solution. Indeed, for 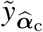, when predicting the bi-modal NF scores, the variability between EEG signals induces decreasing correlation with the reference NF score *y*_c_. Also NF-EEG scores can always be computed from available EEG signals, which leave only NF-fMRI scores to estimate from EEG signals. However, one might want to improve the selection of features for the computation of NF-EEG scores, but this raises different questions about the validation and the reference score. Here we assumed that the given NF scores are relevant to the task and good enough to be considered as reference scores. As expected, for the prediction of the bi-modal NF scores by 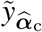, when decomposing the activation pattern 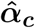 (from 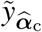), into the 4 matrices corresponding to the different design matrices, the weights corresponding to ***X***_0_(0 second of delay) are mainly located above C3, which is the centre of the Laplacian used for the computation of the NF-EEG scores. This seem to support the fact that with a delay of 0 seconds, only the part of the NF coming from the EEG brings information.

A possibility to increase the prediction of NF-fMRI scores, would be to use more NF sessions as learning sessions, since as observed, EEG signals bring variability in the prediction. Each new bi-modal neurofeedback session could be added to the subject-specific model, to better adapt the model to the subject or patient. This will be investigated in a next study. Given the improved correlation of the proposed NF predictor with bimodal NF scores, it would be interesting in future work to validate its improved performance in actual NF sessions, compared to classical NF-EEG scores. In particular to assess the response of subjects to the predicted bi-modal NF scores and in particular the predicted NF-fMRI scores learned by the proposed model over a standard NF-EEG neurofeedback session, a new and large enough study is needed, as subjects can learn at different pace to regulate their own brain activity.

Presently, the proposed model learns an individual or specific model for each subject that allows a personalised model for adapted neurofeedback sessions. Also a change in strategy for the task (here no specific strategy to imagine moving their right hand was given to the subjects) might impact the learned model, as the relation between EEG and fMRI signals can change. However, in a future work, we are investigating an adaptation of the methodology for the extraction of a common model, taking into account the differences between sessions and subjects, allowing the prediction of NF-EEG-fMRI scores on new subjects who did not participated to the model construction. The model might be less specific, but this would give access to neurofeedback sessions of bi-modal quality using EEG only, for subjects with MRI contraindications, and/or drive a subject-specific model estimation, respecting the strategy used by the subject to progress in its neurofeedback task. The learned NF predictor is of course specific to the particular task considered during the learning session. One can expect to generalise the approach to other tasks where spatial sparsity is relevant. Its extension beyond such tasks is more challenging.

Other ways to improve the method proposed here would be to investigate the use of dynamic functional connectivity, a relatively recent field in BOLD fMRI which needs further investigations to be used along with EEG data (Tagliazucchi and Laufs, 2015). Dynamic functional connectivity would allow to take into account the different network (reward, control and learning) involved during neurofeedback sessions (Sitaram et al., 2017). Dynamic functional connectivity study the temporal fluctuations of the BOLD signal across the brain, and appears to be a promising approach in the EEG-fMRI research field (Allen et al., 2018). However, one should be careful with the potential unknown remaining noise coming from the MRI during EEG-fMRI simultaneous recording, that might unsettle the EEG signal coherence between electrodes.

The long-term objective of our project is to learn from EEG-fMRI NF sessions to provide, outside the MRI scanner, enhanced NF-EEG sessions (Figure 1). A future work will investigate the portability of the learned model (on EEG-fMRI neurofeedback data), outside the MRI scanner, bringing new challenges as dealing with the remaining noises in the MRI after artefact correction, and the absence of ground truth once the EEG is measured outside the MRI scanner.

To conclude, the proposed model here is able to provide a good enough prediction of the NF-fMRI scores, to overcome the absence of NF-fMRI and allows to significantly improve the estimation of NF-EEG-fMRI scores when using EEG only.

## 6 AUTHOR CONTRIBUTIONS

CC guarantor of integrity of entire study. Study concepts and design, data analysis and interpretation, all authors; manuscript drafting or manuscript revision for important intellectual content, all authors; approval of final version of submitted manuscript, all authors.

## 8 ACKNOWLEDGEMENTS

This manuscript has been released as a Pre-Print at (Cury et al., 2019).

Data acquisition was supported by the Neurinfo MRI research facility from the University of Rennes I. Neurinfo is granted by the the European Union (FEDER), the French State, the Brittany Council, Rennes Metropole, Inria, Inserm and the University Hospital of Rennes. This work has received a French government support granted to the CominLabs excellence laboratory and managed by the National Research Agency in the “Investissements d’ Avenir” program under reference ANR-10-LABX-07-01. It was also funded by Brittany region under HEMISFER project.

